# Automated system for training and assessing string-pulling behaviors in rodents

**DOI:** 10.1101/2023.07.02.547431

**Authors:** Gianna A. Jordan, Abhilasha Vishwanath, Gabriel Holguin, Mitchell J. Bartlett, Andrew K. Tapia, Gabriel M. Winter, Morgan R. Sexauer, Carolyn J. Stopera, Torsten Falk, Stephen L. Cowen

## Abstract

String-pulling tasks have been used for centuries to study coordinated bimanual motor behavior and problem solving. String pulling is rapidly learned, ethologically grounded, and has been applied to many species and disease conditions. Typically, training of string-pulling behaviors is achieved through manual shaping and baiting. Furthermore, behavioral assessment of reaching, grasping, and pulling is often performed through labor intensive manual video scoring. No system, to our knowledge, currently exists for the automated shaping and assessment of string-pulling behaviors.

Here we describe the PANDA system (Pulling And Neural Data Analysis), an inexpensive hardware and software system that utilizes a continuous string loop connected to a rotary encoder, feeder, microcontroller, high-speed camera, and analysis software for assessment and training of string-pulling behaviors and synchronization with neural recording data. We demonstrate this system in unimplanted rats and rats implanted with electrodes in motor cortex and hippocampus and show how the PANDA system can be used to assess relationships between paw movements and single-unit and local-field activity. We also found that automating the shaping procedure significantly improved overall performance, with rats regularly pulling >100 meters during a 15-minute session.

In conclusion, the PANDA system will be of general use to researchers investigating motor control, motivation, and motor disorders such as Parkinson’s disease, Huntington’s disease, and stroke. It will also support the investigation of neural mechanisms involved in sensorimotor integration.

**Highlights:**

- High-speed tracking of continuous grasping and pulling behaviors.
- Automated and adaptive reinforcement of string-pulling behavior.
- Integration with neural recording and video tracking systems.
- Open-source software and hardware.

**Supplemental Files:** The supplemental pdf contains additional designs and behavioral data and has been uploaded.

Source code and 3D and laser cut design files can be found at: https://github.com/CowenLab/String_Pulling_System/

Videos are in the GitHub repository at: https://github.com/CowenLab/String_Pulling_System/tree/main/Videos

## Introduction

Skilled reaching movement in a variety of species are studied extensively in the fields of ethology, motor control, motor learning, and movement disorders [1]. The investigation of skilled motor behavior in animal models has advanced our understanding of the neural basis for motor control [2–5], movement disorders such as Parkinson’s Disease [6–9], brain-machine interfaces [10–13], motor recovery following stroke [14,15], spinal cord injury [16], and traumatic brain injury [17,18]. The ability to extend findings from animal models to human behavior is facilitated by the similarities in grasping behaviors in rodents, non-human primates, and human subjects [19]. Several canonical reaching behaviors have been studied extensively in rodents. These include food-pellet grasping tasks [20–23], the vermicelli handling test [24], where animals bimanually manipulate pieces of pasta, the accelerating rotarod test [25] in which rodents balance on a rotating rod, and the string-pulling task that requires bimanual pulling of a string to obtain a reward [26,27].

While each of these tasks is useful, they have limitations. For instance, the rotarod test produces only one behavioral outcome measure (time to fall) which limits the evaluation of fine motor control and the collection of neural data. In contrast, while the single-paw center-out test or skilled pellet-reaching task involve complex grasping behaviors [23,28], these tasks typically involve considerable manual video scoring to identify movements, involve considerable amount of training, and do not require bimanual coordination. These tasks are also compromised when animals prefer a specific limb due to natural preference or due to a lesion/manipulation [17,29]. Limb preference can slow training on the task and complicate interpretation of the data. Bimanual tasks, such as the vermicelli handling test [24] and string-pulling can overcome these limitations. For example, the vermicelli task requires the complex bimanual manipulation of an object; however, assessment of performance requires time-intensive manual segmentation of paw movement [24] and movement time [30], and this is made more difficult by the unconstrained position of the animal relative to the camera or observer.

String-pulling behaviors have received renewed interest as they overcome many of the limitations of the previously described tasks. String-pulling has been used to assess behavior in more than 160 species [1] and have been recently used for the investigation of motor control, movement disorders, and stroke [26,31,32]. In the typical rodent version of this task, baited strings are draped over the walls of the animal’s cage. Animals then make paw-over-paw movements to pull the strings to access rewards tied to the end. This bimanual action is similar to motions naturally performed when pulling nesting material or plants for food [31] and climbing. Perhaps for this reason, rodent versions of this task requires less than a week of training [26]. Despite the usefulness of string-pulling behaviors for basic and translational research, no integrated system currently exists for the automated training and assessment of string-pulling or for the synchronizing of string-pulling data (e.g., paw trajectory and acceleration and string speed) with neural recording data.

Here we describe the PANDA system (Pulling And Neural Data Analysis), an open-source hardware and software system for training, controlling, and analyzing motor behavior and its neural correlates during string-pulling. We also demonstrate the system’s usefulness for characterizing precise bimanual movements in rats (see **Figure 1**) and associating these movements with neural activity (**Figure 4**). The PANDA system allows for measurement of behavioral features such as string speed, paw trajectory, movement phase, and animal posture as well as triggering the automatic food delivery for a specified or algorithmically determined length of string pulled. A key feature of the system is a “continuous loop” (see Crutchfield, 1939) of string connected through a pulley system and attached rotary encoder (**Figure 2**). This design encourages animals to pull longer string distances than traditional procedures (> 100 meters per 15-minute training session) and automatically reinforces animals with liquid food reward for pulling pre-specified distances. Traditional assessment of reaching behavior and paw trajectories involves manual frame-by-frame scoring which is a time intensive process [24,26,31]. The integration of the string-pulling system with a high-speed (≥ 350 frames per second (FPS)) camera allows automated tracking of paw and nose position and the precise segmentation of the reach/grasp movements into specific phases [31]. Finally, we demonstrate how output from the PANDA system is synchronized with neural recording data acquired from the rodent motor cortex and hippocampus to achieve millisecond-level assessment of neural responses to each movement (**Figure 4**). We also show how these recordings can be used to identify neurons that respond to specific reach/grasp phases and how the detailed analysis of the string-pulling behavior could improve assessment of dysfunction in animal models of Parkinson’s Disease (PD). System and analysis code, 3D files for 3D printing, and circuit diagrams are available on GitHub (https://github.com/CowenLab/String_Pulling_System).

**Figure 1.**
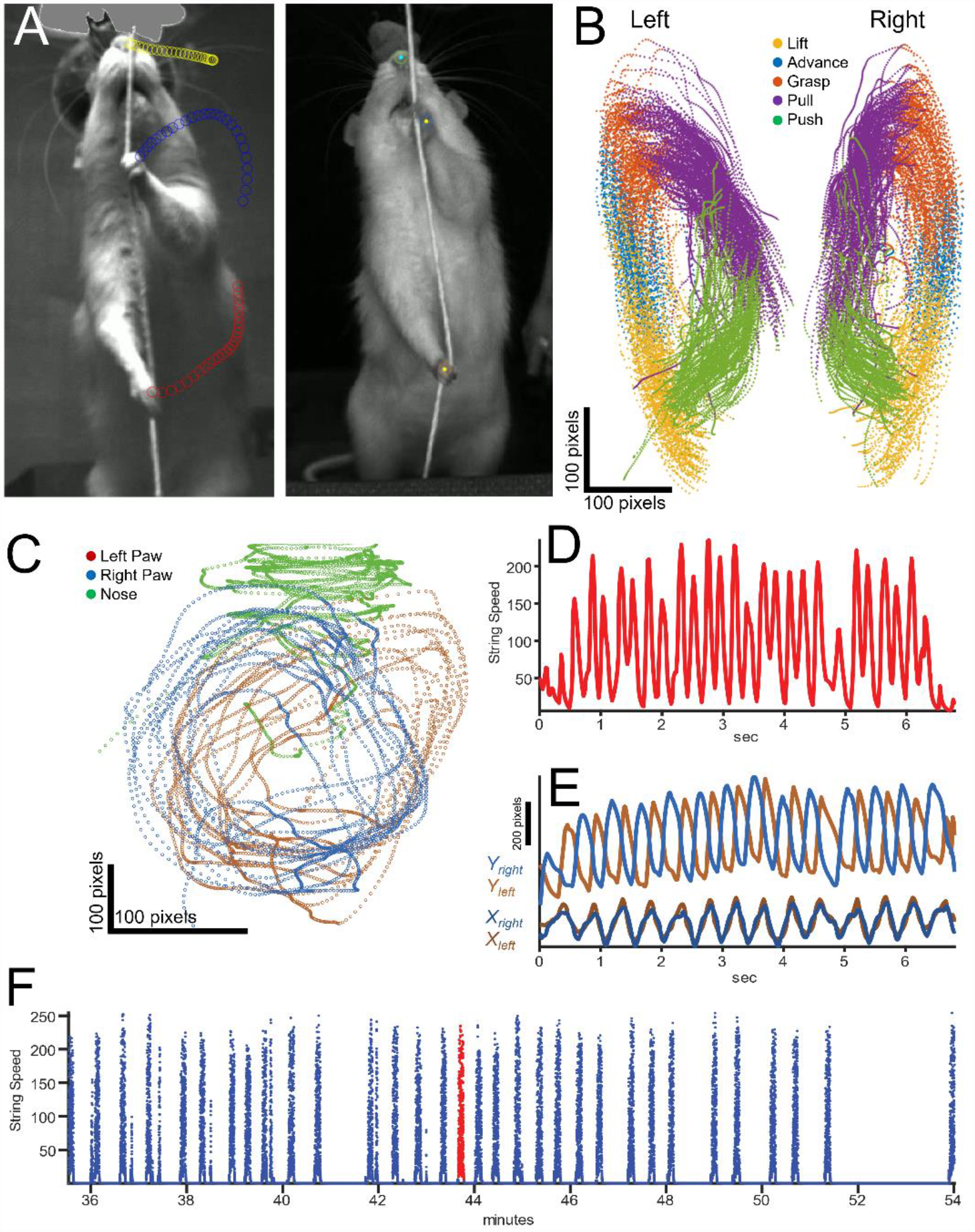
Video Tracking and Behavioral Data: A) Still photos from video recordings of two rats acquired during separate recording sessions. Dots on the left photo indicate automated tracking of the nose and paws using DeepLabCut. B) Paw tracking and automatic segmentation of the 5 phases of string pulling (lift, advance, grasp, push, pull) for a 20-minute behavioral session (33 bouts). Plots for the left and right paw are separated by 100 pixels in the x dimension to improve visualization (to limit overlap). C-E) Trajectories for a single 7s bout of string pulling. C) Paw and nose tracking during a single 7-second pulling bout. X data is expanded relative to Y (see scale bars) to improve visualization of the left and right paw. D) String speed as measured from the rotary encoder during the 7-second bout. E) X and y position of each paw during the 7-second bout. F) String speed through the entire 20-minute training session with each bout indicated as a spike in speed. Red indicates the bout presented in plots C-E.

**Figure 2.**
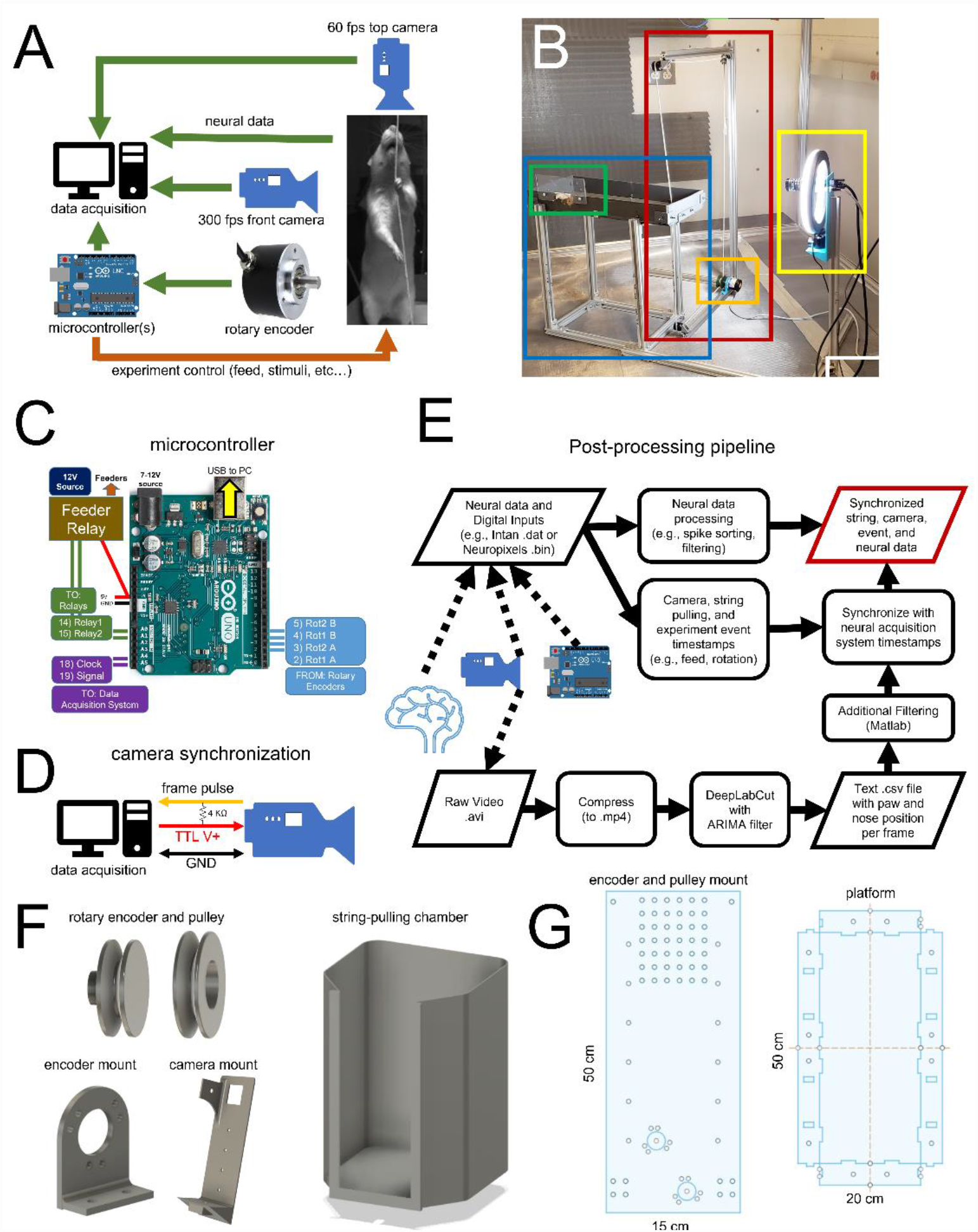
A) Schematic of the data acquisition and experiment control system. Arrows indicate the flow of information from sensors (e.g., camera, rotary encoder, neural signals) and signals for the control of events such as food and cue delivery. B) Photo of the simplest version of the system and components. Red box: Continuous loop of string positioned around a series of pulleys. Orange box: rotary encoder. Blue box: elevated platform. Green box: Food dispenser connected to a solenoid feeder for delivering liquid food (gravity fed, solenoid not shown). Yellow box: High-speed camera with a ring light. C) Connections and pinout for the single-microcontroller version of the system using an Arduino-compatible microcontroller. Key inputs and outputs are indicated. D) Schematic connection with the camera that allows synchronization of each frame with neural/behavioral data. The frame signal and ground are sent to the acquisition system through a BNC cable. The power for the camera is provided by the USB connection. E) Flow chart of the post-processing pipeline of the neural, experiment control, and video data. F) Open-source 3d printed pulleys and wheels used for the string-pulley system, camera mount, and pre-training string-pulling chamber. G) Laser-cut designs for 1) mounting the pulleys and wheels on the extruded aluminum frame, 2) pre-cut base for mounting the microcontrollers and relays in the control box, and 3) the elevated platform.

## Materials and Methods

### System Design Overview

The two main objectives of the PANDA system are 1) to facilitate precise assessment of bimanual behaviors and 2) to quantify their neurophysiological correlates. A photo schematic and images of key components of the system are shown in **Figure 2ab**. Here we describe the application of the PANDA system for monitoring a single string/rotary encoder; however, the described system also accepts input from a second rotary encoder for a dual-string setup (*e.g*., for 2-choice behaviors). Components of the system and manufacturers are listed in **Table 1**. A table with full website links is also provided in **Supplementary Table 1**. Major components include an elevated platform for the rat with a solenoid-controlled feeder at one end and the string apparatus at the other. The string apparatus consists of a loop of cotton string attached to a pulley system with one pulley being connected to a rotary encoder for measurement of string speed and direction. An Arduino-compatible microcontroller tracks string speed and pulled distance and triggers liquid reward (Ensure™) delivery by activating a solenoid valve. Outputs from the microcontroller encode events such as string speed and food delivery and send these signals to an attached PC (via USB) and/or a neural data acquisition system. For the experiments reported here, we used the Intan neural recording system (Intan Technologies Inc.), but any system that accepts digital or analog input will work. A high-speed camera (> 350 FPS, Allied Vision Inc.) collects video data of body, head, and paw movement which is processed off-line using DeepLabCut [33]. Software for controlling the PANDA system and for processing acquired video and string-pulling data is described below and available at https://github.com/CowenLab/String_Pulling_System.

**Table 1.** Parts and Suppliers. System cost without the neural recording system is approximately $1200.

| **Table 1 - Parts List** |  |  |
| --- | --- | --- |
| **Component** | **Manufacturer** | **Part No.** |
| **Electronic Components** |  |  |
| Mako U-130b Camera* (option 1) | Allied vision | Mako U-130b |
| Alvium 1800 U-040 (option 2) | Allied vision | Alvium 1800 U-040 |
| LED Panel | Viltrox | L116T |
| Ring Light, 10" | UBeeSize | UBeeSize Ring Light |
| Arduino Uno (RRID:SCR_017284) | Arduino | Arduino Uno Rev3 |
| Rotary Encoder | BQLZR | BQLZR 600P/R |
| Digital Distance Sensor 5cm | Pololu | 4050 |
| Project Box 12.2 x 11.2 x 4.5" | Zulkit | NA |
| Solenoid Valve 1/4" | STC Valve | 2P025-1/4 |
| Panel Mount Aviation Connectors | Hilitchi | 8541770567 |
| **Physical Apparatus** |  |  |
| T-slot nuts | Sutemribor | STBR-T-luomu-160P-kit |
| Cotton Twine #16 [CHECK] | Ace Hardware | C8016B0008AC |
| 3D printer wheel with bearings | SeekLiny | NA |
| Extruded Aluminum (20/20) | Zyltech | EXT-2020-REG-1000-10X |
| 90° Corner Bracket | LANIAKEA | NA |
| **Neural Data Acquisition Systems** |  |  |
| Intan USB Interface Board (option 1)* | Intan Technologies |  |
| RHD Recording Controller (option 2) | Intan Technologies |  |
| Neuropixels recording system | Imec |  |
| *\* Model Discontinued. The Alvium 1800 is the updated model.* |  |  |

The list does not include common parts such as nuts and bolts, wire, header connectors, standoffs, etc...

#### Monitoring Speed and Distance Pulled with a Rotary Encoder

String pulling speed and distance were detected using a two-phase rotary encoder with a resolution of 600 pulses per rotation (see **Table 1** for supplier and model). Signals from the encoder were processed by an Arduino-compatible microcontroller to provide real-time measures of rotation speed, direction (up or down), and string length pulled. To reduce computational demands on the Arduino and the data acquisition system, incoming pulses from the rotary encoder were downsampled from 600 to 30 pulses (tics) per rotation resulting in a resolution of 18 degrees or ∼0.34 cm of string pulled per tic. Initial calibration of the number of tics per centimeter of string pulled was determined by manually pulling the string a known distance and measuring the number of tics. Calibration only needs to be performed once.

This single-microcontroller system was quite effective for shaping and assessing simple string-pulling behaviors such as rewarding animals for a given length of string pulled. We found that a dual-microcontroller setup was more practical for more complex experiment control scenarios involving additional inputs, effectors, and complex contingencies. In this variant, one controller was dedicated to monitoring the rotary encoder and another was dedicated to experiment control (*e.g*., processing inputs, calculating reward contingencies, triggering effectors, etc). The wiring for the dual-microcontroller version of the system is presented in **Supplementary Figure 1**.

The entire system can operate autonomously without requiring a connected PC or data acquisition system. However, for synchronization and logging, digital and USB serial output from the microcontroller indicating rotary encoder rotation, direction, distance pulled, and events such as the delivery of reward can be sent to a data collection PC via serial output (USB) and/or to a neural data acquisition system via 5V TTL pulses (see **Data Acquisition and Synchronization with Neural Data** below).

#### String System Hardware Design and Construction

The continuous loop of string was made by looping the string over the rotary encoder and 3 pulleys mounted on the corners of a C-shaped adjustable frame (**Figure 2b**). The frame was made of 2020 aluminum (Zyltech Houston, Texas, United States, see **Figure 2b**). Each pulley used low-friction steel-bearing wheels used in most 3D printers (**Table 1**). Wheels were either embedded in a 3D printed shell in the shape of a pulley or two wheels were sandwiched together with a nut and bolt to form a pulley (see **Figure 2b** and **2f**). The continuous loop of string was wrapped once around (360 degrees) the wheel connected to the rotary encoder to reduce slipping and increase accuracy. The C-shaped aluminum frame housing the pulley system was positioned at one end of the elevated rectangular platform (20 cm x 50 cm). The top of the frame was at least 40 cm above the platform so that it could not be grasped by the rat and would not interfere with video recording. This platform was constructed from painted wood and extruded aluminum and had a liquid food reward port at one end. An elevated platform was used as it discouraged animals from jumping off the apparatus, allowed animals to easily view allocentric cues, and eliminated opportunities for the animal to hit their neural implant against a wall. Designs for a laser-cut version of the elevated platform (**Figure 2g**) are provided in the GitHub repository. While the elevated platform was optimized for neural recording, we also developed a fully enclosed and non-elevated training chamber that was useful for pre-training (**Figure 2f right**). Designs for the FDM-printed and fully enclosed chamber are provided on the GitHub site.

#### Software and Hardware used for 3D Design and Manufacturing

3D printed parts such as pulleys and mounts were manufactured in-house using a Creality CR-10S FDM printer (Creality Inc., https://creality3d.shop/). Laser cut parts were manufactured using a Sculpfun S9 laser cutter (Sculpfun Inc., https://www.sculpfun3d.com/). All 3D (.stl) and laser cut (.dxf) designs and files are available in the GitHub repository. 3D printing and laser cut files were created using SolidWorks (https://solidworks.com) or Autodesk Fusion 360 (https://autodesk.com).

#### Video Monitoring of Behavior

A high-speed video camera was mounted on a 2020 aluminum frame and placed 60 cm in front of (facing) the rat (**Figure 2b**). A second camera was mounted 80 cm above the rat to monitor the location of the animal. Each camera sent a digital pulse to the acquisition system at the start of each frame allowing frame-by-frame synchronization of behavioral and neural data. Video from the string-facing camera was captured using a Mako monochrome U-130b camera (Allied Vision, Stadtroda, Germany) configured using the Image Acquisition Toolbox in Matlab 2019b or StreamPix Lite software (NorPix Inc, Montreal, Canada). It should be noted that Allied Vision has recently replaced the Mako camera with the Alvium 1800 U-040, and this camera has similar or superior features to the Mako. A frame rate of 367 FPS was achieved using the Mako camera at 600 × 850-pixel resolution (grayscale). This high sampling rate increased tracking precision and reduced motion blur during fast grasping motions relative to standard 30 or 60 FPS USB cameras. The quality of the video was improved by using a ring light (UBeeSize, City of Industry, California, United States, see **Figure 2b**). The camera and ring light were secured to the 2020 aluminum arm with a custom 3D printed mount (**Figure 2d**).

#### Data Acquisition and Synchronization with Neural Data

Output from the string-pulling system microcontroller is sent to 1) an acquisition computer via USB and 2) to a neural data acquisition system through digital TTL signals. Data sent via USB indicating experimental events and data was also recorded as a comma-separated-value file on the host PC by using a common serial communication program (Putty, https://putty.org/). These data included rotary encoder movement (resolution of 1/20^th^ of a turn or 18 degrees), the direction of the rotation, and events such as the time of food delivery, and beam-breaks from laser proximity sensory (Pololu Inc.). Digital pulses relaying this same information were also sent to the neural acquisition system.

#### Post-Processing Collected Video Data

Once data was collected in the formats described above, it was post-processed according to procedures summarized in **Figure 2e**. Video was recorded as uncompressed .avi files and then compressed to .mp4 format using Format Factory (Free Time, Inc.). Video recording was triggered shortly after the data acquisition system began recording data to ensure that the first frame was logged. Video files were processed using DeepLabCut [33] to extract paw and nose position in each frame (described below). Frames were selected from the raw video (typically 108 frames) and manually scored to identify the paw and the nose. This scored data formed the training and testing sets, and DeepLabCut was trained for ∼730,000 iterations on an 85:15 train:test split until loss converged to 0.003. Analysis with the network produced CSV files containing the positions of the tracked paws and nose as well as a confidence measure for each position estimate. These data were filtered further in DeepLabCut using an autoregressive integrated moving average (ARIMA) filter (AR=3 and MA=1). These data were loaded into Matlab for the extraction of additional features (e.g., paw speed, angle between the paws, position relative to the nose, and automatically segment each pull into lift, advance, grasp, pull, and push phases).

#### Validation of Paw Tracking

DeepLabCut provides validation metrics such as distance in pixels of the estimated feature positions from the manually labeled positions in the training and testing sets of images. Analyses of these data showed a 1.04-pixel (∼0.027 cm) error for the training split images and a 4.33-pixel (∼0.105 cm) error for the test images. For our video recording configuration, rat paws had widths of ∼35 pixels or ∼0.92 cm, which was approximately 9-fold wider than our test error (4.33 pixels, ∼0.105 cm).

#### Behavioral Segmentation

Each pull of the string was automatically segmented into reach and withdrawal phases as well as more fine-grained “lift, advance, grasp, pull, and push” phases. These phases were comparable to those described in Blackwell *et al*., [34] (See **Figure 1b, 3cd)**. This was accomplished through a three-step process. First, the time series of vertical (y) paw data was smoothed (Matlab: movmean() with a 4 second window) and band-pass filtered (1-8 Hz). The Hilbert transform of this output yielded a continuous measure of pull phase (Matlab: angle(hilbert(y_data))). Second, the velocity (positive up, negative down) and acceleration were determined from the x and y time-series data. Third, data from each paw was analyzed cycle-by-cycle to identify the categorical label of each pull phase (e.g., lift, advance, grasp, pull). For example, the transition from “pull” to “push” was determined as the time when the paw position during a downward motion (negative velocity) reached its maximum eccentricity in the x dimension. Segmentation accuracy of the automated procedure was confirmed visually by comparing videos of each reach or withdrawal to the output of the automated procedure.

### Training the String-Pulling Behavior

Prior to training, rats were handled for 10-30 minutes per day for 1-2 weeks to accustom them to experimenters. At the onset of training, rats were food-restricted according to the Institutional Animal Care and Use Committee (IACUC) guidelines and weighed after each training session. Food was provided to ensure that animals remained ≥ 85% of their free-feeding weight. Training the string-pulling behavior was modeled on previously published procedures [26] but modified to allow habituation of the rats to the elevated platform and to reinforcement through the solenoid-driven liquid food feeder (Ensure™). Animals were trained in the string-pulling behavior in 3 phases with the performance criterion being that each animal achieve ≥ 20 bouts of string pulling of ≥ 1 meter per bout during a single 20-minute training session. All rats described here reached this criterion in 7-14 days. While this was the minimum threshold, many animals exceeded this level of performance with one animal pulling > 700 meters in a 1-hour session.

#### Phase 1 – pull in home cage

As in [26], animals were habituated in a walled arena or home cage with 20 strings of lengths 30-100 cm draped over the edge. Half of these strings were baited with a Cheerio. The rats were allowed to interact with and pull the string to receive the reward. Initially, animals were encouraged to pull by receiving a half Cheerio for a partial pull of the string. Rats were left in the arena for one hour or until all strings were pulled inside the cage. This continued for two sessions. During Phase 1, rats were also placed on the string-pulling platform for 5-10 minutes/day without the string or food reward so that they would become accustomed to the elevated platform. During Phase 1, a small amount of Ensure was placed in each animal’s home cage to habituate them to the reward.

#### Phase 2 – manual reinforcement on platform

Rats were placed on the elevated string-pulling platform and rewarded for investigating the string by hand-feeding with Cheerios. Animals quickly began pulling the vertical string for short bouts and were hand-fed each time 1-meter of string was pulled. The length required for reinforcement was gradually increased to 3 meters until this length was pulled 3-4 times during a 30-minute training session. This performance was typically achieved in 1 day at which point training moved to Phase 3.

#### Phase 3 – automated shaping

Rats were placed on the elevated platform and were rewarded with Ensure (via solenoid) delivered to a dish 120 cm from the string. Shaping and reward delivery was controlled by the microcontroller where reward was initially delivered for any movement of the string and then the criterion distance was gradually as animals consistently pulled each target distance (e.g., 5cm, 20 cm, 50cm, 100cm) until the criterion for that day was met.

### Animals

Data collected from n = 14 rats are reported here from a variety of experiments that demonstrate the utility of the system under various experimental conditions. All animals were male, Sprague-Dawley rats (∼275-350 g at time of arrival; Envigo RMC Inc., Indianapolis, IN) and were single housed in a temperature and humidity-controlled room on a 12-hr reverse light/dark cycle. Food and water were provided *ad libitum* for the duration of the habituation period. When training began the rats were food restricted to 85% of their body weight. All animal procedures were in accordance with University of Arizona IACUC and federal NIH guidelines for the Care and Use of Laboratory Animals. Animals were divided into the following groups: **Single-Unit Recording in M1 and striatum** (n=2), **Local-field Recording in the Dorsal Hippocampus** (n = 6), **Naïve Non-implanted rats** (n = 6) for assessment of learning, and an **Animal model of Parkinson’s disease** (n = 3 for investigator-scored experiment and n = 1 for the automated string pulling apparatus experiment) and **sham-lesioned animals** (n = 3).

### Stereotaxic Surgeries for Rats Implanted with Chronic Microdrives

On the day of surgery, rats were anesthetized using 1.0 - 2.0% isoflurane in oxygen (flow rate 1.5 L/min) and placed into a stereotaxic apparatus. The microdrive was centered over a craniotomy made on the right hemisphere (Coordinates relative to bregma: Hippocampal experiments: ML: 2.0, AP −3.8 mm. M1 experiments: ML: 2.2 AP: 1.5mm). Tetrodes were constructed of four twisted polyimide-coated nichrome wires (13 μm diameter). General procedures for microdrive surgeries are described in [35,36].

### Stereotaxic Surgeries for the Rat Parkinson’s disease model

The unilateral 6-hydroxydopamine (6-OHDA) lesions to model PD were done as published in [37]. The lesions were done with 6-OHDA injection in the medial forebrain bundle, the coordinates for the 6-OHDA injection relative to bregma: AP = −1.8 mm, ML = +2.0 mm, DV = −8.2 mm and AP= −2.8 mm, ML = +1.8 mm, DV = −8.2 mm. Mean amphetamine-induced rotations of the animals included in the manual study were performed at 3-weeks post-lesion: 13.9 ± 5.1 SEM, indicating a >90% lesion.

### Data Analysis

All post-processing, filtering, and statistical analyses were performed using Matlab2019b and Python 3.6. Statistical analyses were performed using Matlab or R with alpha = 0.05. Unless otherwise stated, the Holm–Bonferroni method was used for multiple comparisons corrections.

## Results

The performance of the PANDA system with implanted and unimplanted rats was evaluated with animals trained as described under **Methods**: **Training the String-Pulling Behavior**. For most analyses, data was analyzed from 6 rats trained prior to microdrive implantation with the objective of each rat pulling an average of 2 meters of string per bout for > 10 bouts during a single 30-minute training session. A bout was defined as a period where the rotary encoder detected motion of the string for ≥ 1 second, and a bout ended when the rotary encoder was not moving for ≥ 1 second. The 6 rats described in **Figure 3b** required < 8 days to meet the 2-meters per bout goal. This level of behavior was maintained following implantation of chronic electrode arrays. These animals completed an average of 42 bouts/session, with individual bouts lasting ∼7 seconds (**Figure 3b**). All behavioral metrics were calculated automatically from the data acquired from the rotary encoder, and thus did not require labor-intensive video scoring.

**Figure 3.**
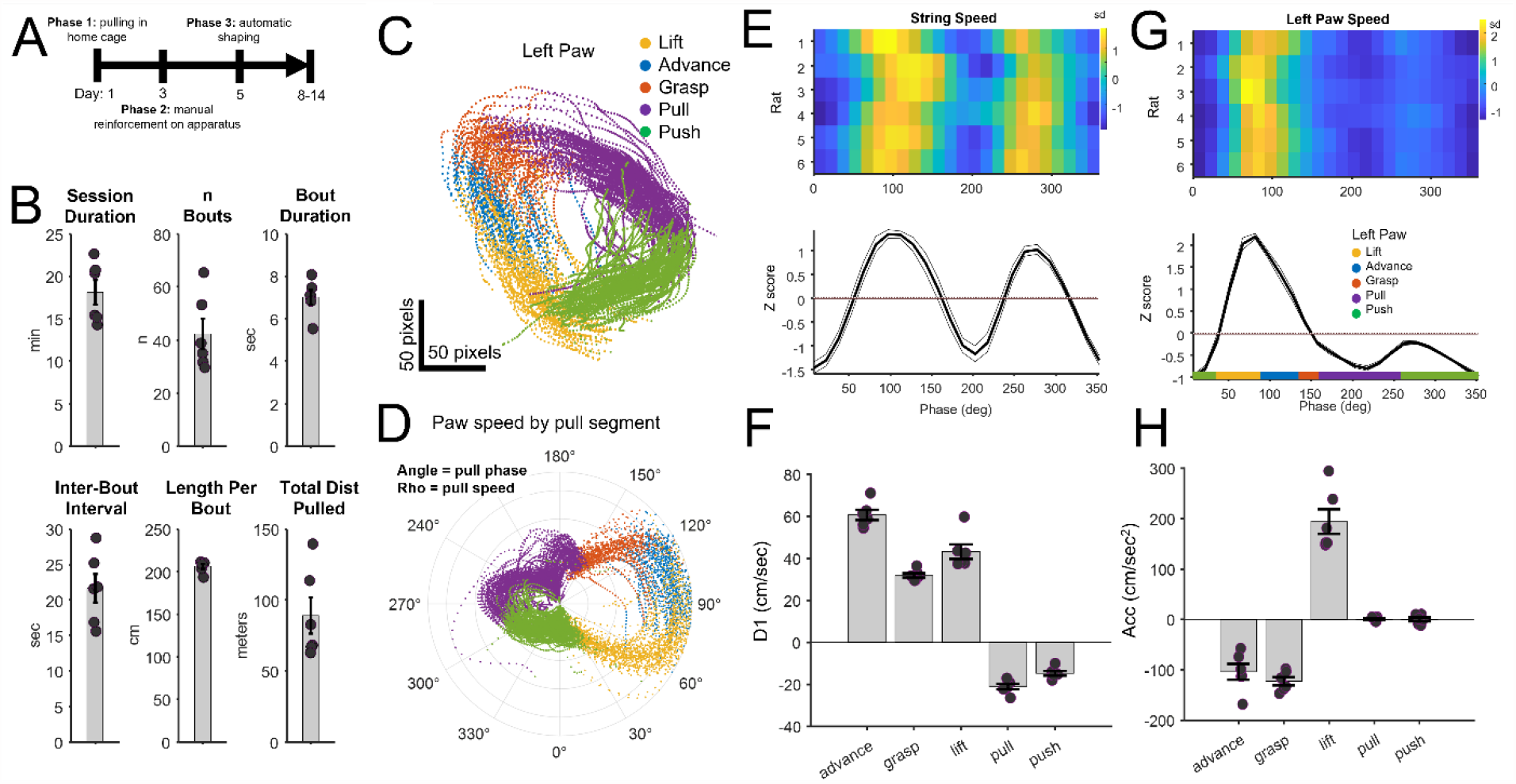
**A)** Timeline for training rats on the string-pulling task. **B)** Behavioral performance of n = 6 rats after 10 days of training. Session duration was the time animals actively performed the task; n Bouts indicated the mean number of pulling bouts per session; Bout duration was the mean duration of each bout; Inter-bout interval indicates the time from the end of one bout to the start of the subsequent bout; Length per bout was the mean length of string pulled per bout; Total Dist Pulled was the total length of string pulled per session. **C-H)** Paw Kinematics: **C)** Paw position by automatically segmented pull phase from a single session (left paw). Data from **Figure 1b. D)** The polar plot indicates paw angle (angle) and pull speed (radius) for each segmented pull phase (color) for the session. The rightward asymmetry indicates the large increase in speed during the lift and advance phases. **E)** Paw velocity (D1 = first derivative of the y coordinate of the paw) for each phase of a pull cycle (n = 6 rats, mean +/-SEM). **F)** Paw velocity averaged per reach/pull phase. Consistent differences in D1 (positive = upward movement, negative = downward movement) were observed for each phase (p < 0.000001, One-way ANOVA, n = 6 rats) and were notably consistent between animals. **G-H)** As with **E-F**, except for acceleration (cm/sec^2^). Acceleration differed between each phase (p < 0.00001, One-way ANOVA, n = 6).

### Paw Kinematics

Kinematic measures included nose and paw position, paw speed, acceleration, and the angle between paws. These measures were calculated using DeepLabCut and through analyses of DeepLabCut output in Matlab. The vertical position of the paw along with paw velocity (illustrated in **Figure 3c**) were used to determine the phase (angle) of the paw through each reach/withdraw cycle (**Figure 3d**). This was accomplished by performing a Hilbert transform and extracting phase from the y-data time series (*e.g*., **Figure 1e**, Matlab: angle(hilbert(y_time_series))). The start of each cycle was defined as the time when phase reached the peak of the y coordinate within each cycle. Visual inspection confirmed that this point corresponded to a time shortly preceding the animal grasping the string. Paw position (x, y, velocity, paw angle) information was used to automatically segment each pull cycle into discrete categories such as reach and withdrawal and more fine-grained categories such as advance, grasp, lift, pull, and push as described in **Methods**. Code segmenting pull cycles is provided on GitHub (Matlab: PANDA_segment_reach_and_withdrawal()).

Consistent measures of paw movement and speed were acquired in all 6 animals (**Figure 3e-h**). **Figure 3e** shows mean string speed for each of the 6 rats as measured by the rotary encoder and aligned to the pull phase of the left paw. A more targeted analysis of the left paw alone (data from the high-speed camera + DeepLabCut) is shown in **Figure 3g**. These data indicate that paw speed was notably fastest during the lift and advance phases. Speed is presented as z scores for visualization purposes as it normalizes inter-animal differences in mean movement speed. Mean movement velocity (speed and direction) in the y dimension (cm/sec) and acceleration (cm/sec^2^) of the left paw for each animal are presented in **Figure 3fh** further demonstrating that each phase of the reach/pull movement is characterized by a unique kinematic profile.

### Neural Responses to String Pulling

The capacity of the PANDA system to track paw, nose, and body movement with millisecond precision suggests applications for the investigation of the neural mechanisms underlying motor control, sensorimotor integration, and movement disorders. Here, we summarize data from the string-pulling system acquired from rats implanted with single-unit and local-field electrodes. To our knowledge, this is the first study to measure single-unit or local-field data during a string-pulling behavior. **Figure 4a** presents motor cortex (M1) single-unit activity (blue dots) overlaid on paw trajectories during the reach and withdrawal phase. In this example, the M1 neuron appears to respond selectively during the withdrawal phase. **Figure 4b** presents data from a second neuron with activity aligned to the start of the push phase. These data indicate that the neuron was strongly modulated by the pull phase with activity peaking shortly before the push phase.

**Figure 4.**
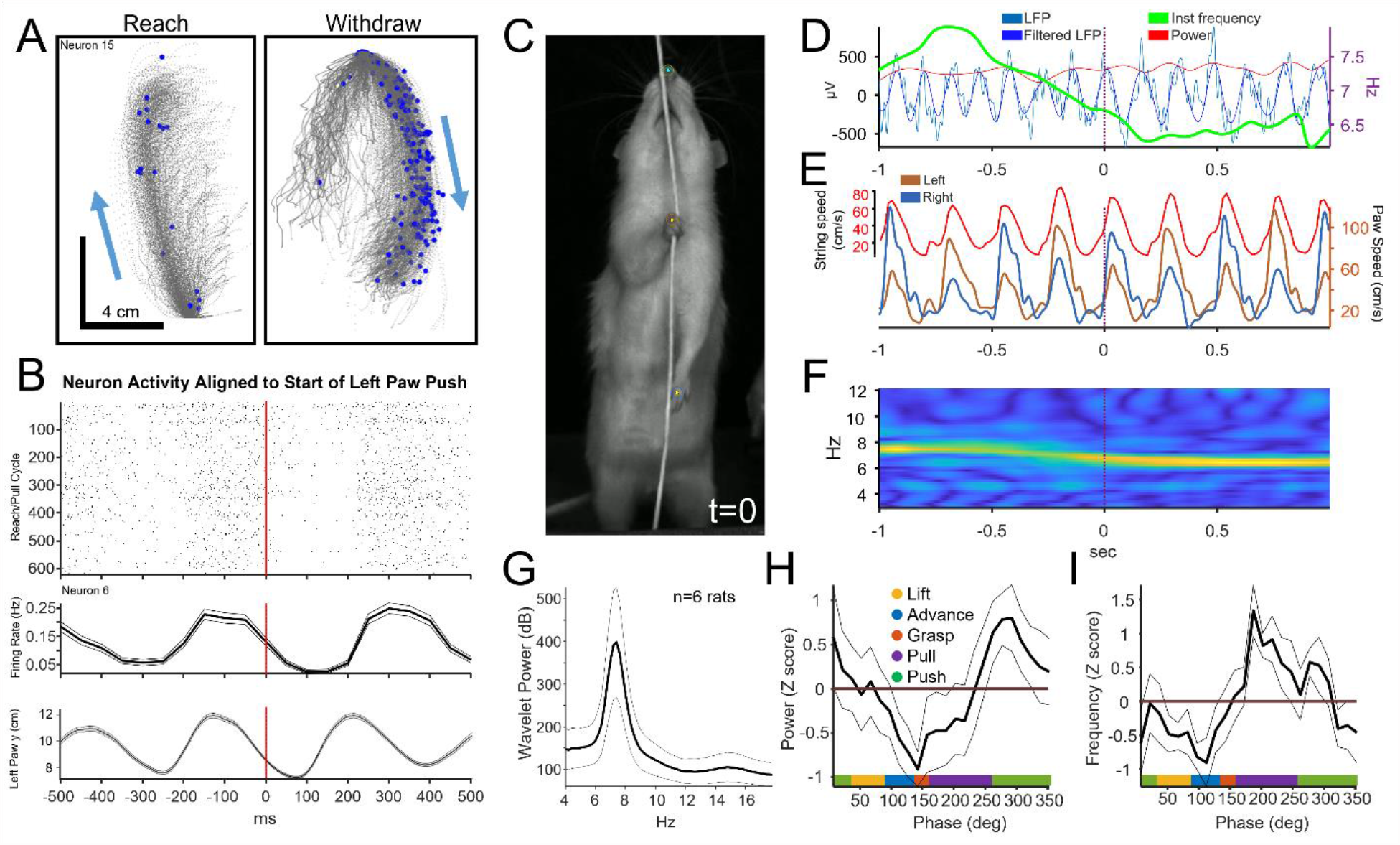
A) An example of an M1 neuron with a selective response to the withdrawal phase during string pulling. Action potentials are indicated as blue dots. B) Rastergram of the activity of another M1 with the timing of action potentials aligned to the start of the push phase. Top: rastergram with each row indicating a single string-pulling cycle. Middle: mean response (+/-SEM). Bottom: y position of left paw. C-F) Local field responses in the theta band (4-12 Hz) aligned to the time indicated by the video frame in C. D) Original and band-pass filtered (4-12 Hz) local field signal along with an instantaneous measure of power and frequency aligned to the frame indicate in C. E) Paw motion (from video) and string speed (from rotary encoder). F) Wavelet spectrogram indicating power in the theta band. G) Mean power spectral density across 6 rats indicating clear theta-band activity during string pulling. Data was restricted to times when the rat was actively pulling the string. H) Mean (+/-SEM, n = 6 rats) theta power measured as a function of the phase angle and segmented phase label of the left paw. This analysis indicated that theta in rats was modulated by the phase of the string-pulling behavior with theta power peaking during the push phase. I) As in H, but for the instantaneous measure of theta frequency. These data indicate that theta frequency peaked near the start of the pull phase.

In a second set of experiments, local-field recordings were acquired from the dorsal hippocampus of 6 rats. In this study, neural responses to theta-band (4-12 Hz) oscillations were examined during the string-pulling behavior. Theta in the hippocampus of rodents is of interest as it is associated with memory formation and retrieval, sensorimotor integration, and spatial navigation [38,39]. There is considerable evidence that theta is impacted by animal motion and incoming sensorimotor information as theta power and frequency are modulated by running speed and acceleration [40–44]. Far less is known about the relationship between theta and the movement of individual limbs or during complex multi-phase behaviors such as grasping/pulling. **Figure 4c-f** presents an example of local-field and kinematic measures acquired from a single rat. All data is aligned to the time of the video frame shown in **Figure 4c**. These data show clear theta-band activity during string pulling (**Figure 4df**). Kinematic data (string speed and paw position) are presented in **Figure 4e**.

Mean spectral response averaged across the 6 rats indicated clear theta-band activity during string pulling (**Figure 4g**). Theta power and frequency were also analyzed as a function of the phase angle and phase label of the left paw (**Figure 4hi**) with the left paw being contralateral to the side of the implant. This analysis indicated that theta in rats was modulated by the phase of the reach/withdraw behavior, with theta power peaking during the push (**Figure 4h**) and frequency peaking near the start of the pull phase (**Figure 4i**). A video showing the aligned neural response to string pulling phase for an individual rat can be found in the GitHub repository.

### String-pulling Behavior in the 6-OHDA Model of Parkinson’s Disease

To demonstrate that string-pulling behaviors have potential applications for translational research in Parkinson’s disease, we conducted experiments in unilateral 6-OHDA-lesioned rats [45]. Two experiments were performed. The first involved pre-training rats on the home-cage string-pulling behavior described in Blackwell *et al*. [34]. A subset of these animals was given 6-OHDA lesions and subsequently evaluated on the string-pulling behavior by blinded investigators. In the second experiment, a detailed kinematic analysis of reaching/pulling was performed using the automated string-pulling system described here. Lesions in all 6-OHDA animals were histologically verified to be >90% and 6-OHDA animals exhibited amphetamine-induced rotations (mean rotation count of 13.9 ± 5.1).

In the first experiment, n = 6 rats were trained using the methods described in Blackwell *et al*. [34] (in the home cage with baited strings draped over the walls, not on the elevated platform). After reaching criterion, n = 3 rats received 6-OHDA lesions and n = 3 received sham lesions. The mean time to complete the task (pull all 10 strings) and the number of missed grasps of the string by the contra-lesioned paw was significantly higher in the 6-OHDA group (**Supplementary Figure 2**). Semi-quantitative western verification of the 6-OHDA lesion for these animals is provided in **Supplementary Figure 3**.

In the second experiment, a detailed analysis of paw kinematics of a single 6-OHDA lesioned animal was performed using the string-pulling system described here (elevated platform, high-speed camera). Kinematic data and histological verification through tyrosine hydroxylase immunoreactivity in the striatum are provided in **Supplementary Figure 4**. Unlike the previous experiment, this rat was only trained to perform the string-pulling task after the 6-OHDA lesion, indicating that lesioned animals can *de novo* acquire the behavior. Right vs. left differences in paw kinematics were observed in this animal during the grasp, pull, and push segments (**Supplementary Figure 4a**). The speed of the paw associated with the lesioned hemisphere (left paw) was faster than the right paw these phases (p < 0.05, Wilcoxson test, Holm correction). This was unexpected as Parkinson’s disease is associated with motor slowing. That said, this single example is primarily illustrative as it lacks a pre-lesion measure that would control for an inherent side preference. Even so, these data indicate that PD-associated alterations in performance could be identified in this behavior and that PD-model animals can learn and perform this task. Future work could investigate whether dopamine loss is associated with aberrant grasp/clasp behaviors as has been shown following motor cortex lesions [46].

## Discussion

Forms of string-, grass-, or rope-pulling behaviors have been observed in >160 species [1,47]. These complex, yet rapidly learned behaviors allow in-depth assessment of bimanual motor dynamics and cognition. Interest in string pulling has grown given their utility for investigating motor function and the behavioral and neural consequences of motor disease and stroke [17,32]. Training and scoring animal behavior on string pulling tasks is typically performed manually. Here we describe PANDA, a hardware and software system for the automated training and assessment of string pulling with a level of precision that supports the analysis of the relationships between fine motor movements and single-neuron and local-field activity. This level of precision is achieved by using a high-speed (>350 FPS) camera, rotary encoder, post-processing with DeepLabCut [33] and custom open-source code for tracking limb motion and neural activity.

### Applications for the Training and Assessment of Motor Behaviors

The integration of a continuous “infinite” loop of string with a rotary encoder allows for precise determination of the length of string pulled and reliable reinforcement. Furthermore, the ability to optimize the reward delivered per pull bout allowed for longer bouts and behavioral sessions. To illustrate, one rat pulled individual bouts of up to 16 meters and a total of 713 meters in a single one-hour session. This behavior provides an unprecedented amount of data for assessment of movement and neural activity. Existing behaviors used for assessing reaching/grasping behaviors, such as the vermicelli grasping task [24] and food-pellet grasping tasks [20–23], do not allow extended training periods nor do they readily support full-body tracking as the animal is typically free to change their orientation relative to the camera.

There are potential applications of the PANDA system for investigating motor disorders such as Parkinson’s and Huntington’s disease, amyotrophic lateral sclerosis (ALS), and stroke given the precision, quality, and quantity of bimanual reaching and grasping data generated. To illustrate, recent research using a manually scored (not automated) string-pulling behavior identified potential motor consequences of stroke and motor-cortex devascularization [32]. Specifically, this study identified unique deficits in grasping and supination/pronation of the paw following devascularization. The automation of the collection of such data and its synchronization to neural activity as described here could significantly advance such research by allowing the investigation of how these behavioral effects are mirrored in the neural activity of motor circuits throughout the brain.

String-pulling behaviors are rapidly learned with animals requiring 7-8 days to reliably pull for ∼40 bouts within a 20-minute session. The rapid learning of this behavior supports between-group assessment of motor and skill learning. For example, the acquisition of the task could be evaluated in animal models of Parkinson’s disease, aging, and Alzheimer’s disease and evaluate the effects of pharmacological treatment. Most behavioral and neuroscience laboratories performing such research could implement the string-pulling system described here with minimal cost as it uses off-the-shelf components and only requires a moderate understanding electronics and coding to build and operate.

### Applications for Assessment of Neural Activity

The string-pulling system was synchronized with the video and data acquisition systems to allow for millisecond-precision analysis of neural and motor activity. To demonstrate this capability, single-unit activity in motor cortex was mapped onto the start and end of the reach phases of the pulling behavior (**Figure 4ab**). In addition, we identified changes in hippocampal theta band activity that correlated with string pulling and with specific segments of the reach/pull motion (**Figure 4hi**). This level of detail in the analysis of motion could help resolve persistent debates regarding the extent sensorimotor and proprioceptive information drive activity within dorsal hippocampus [48]. In a different domain, it is conceivable that large datasets of synchronized neural and grasping/reach data on string-pulling task could support development of improved algorithms for brain-machine interfaces (BMI).

### Limitations

While video data collected from our system integrates well with DeepLabCut, it is not currently compatible with a recently developed Matlab-based approach for analyzing string-pulling behavior developed by Inayat *et al*. [49]. This is largely due to the use of high-speed cameras that only collect data in grayscale, where the Inayat *et al*. [49] software requires color video data for segmenting paw and string position. It is conceivable that future versions of our system could use a different camera and incorporate color video; however, this would likely come at the cost of reduced sampling rates.

Improvements could also be made in the precise segmentation of the reach and grasp motion. For example, using visual scoring, Blackwell *et al*. [26] observed subtle changes in elbow position during different phases of the reach/grasp motion. It is conceivable that, with additional hand-scored training data, DeepLabCut could be trained to track elbow position and the angle between the elbow and the paw as well as the position of each paw digit for assessment of grasping. Such data could allow more fine-grained automated segmentation of movement.

## CRediT authorship contribution statement

Cowen: Design of the system, software development, writing manuscript, analyzing data, building the system.

Jordan: Hardware development and 3D designs, control software development, writing manuscript, analyzing data.

Falk: Revising manuscript, supplying, and advising on use of 6-OHDA animals. Bartlett: Histology, surgical procedures, and work with 6-OHDA animals.

Sexauer: Training 6-OHDA animals, assessing behavioral data. Stopera: Western blot validation of 6-OHDA model animals.

Tapia: Building and testing the string-pulling apparatus.

Winter: Implementing the Matlab-based analysis of video described in Inayat et.al. 2020.

Vishwanath: Training animals, assessing behavioral data, recording neurophysiological data.

Holguin: Training animals, assessing behavioral data, recording neurophysiological data.

## Data availability

Data, 3D designs, and software is available on GitHub at https://github.com/CowenLab/String_Pulling_System.

## Acknowledgments

SC: National Institute of Health R01 NS123424, Evelyn F. McKnight Brain Institute

TF: National Institute of Health R56 NS109608 and R01 NS122805

