## Supplementary Figure for "Automated system for training and assessing string-pulling behaviors in rodents"

### Dual Microcontroller Option

In this option, a dedicated Arduino-compatible “Rotary Encoder Microcontroller” (REM) is used to pre-process the signals from the rotary encoder. The processed data is then sent to an “Experiment Control Microcontroller” (ECM) and to the data acquisition system. The REM down sampled the signal from 600 to 30 pulses/tics per rotation (18 degrees or ~0.34 cm of string pulled per tic) and determined rotation direction (clockwise or counterclockwise). Down sampling was useful as 600 pulses per rotation strained the data acquisition system’s digital input tracking capabilities when animals pulled at full speed. Furthermore, while it was possible to perform simple experiment control functions such as monitoring behavior and delivering rewards using a single microcontroller, we found that separating these components into separate microcontrollers freed up computational resources on the ECM and allowed easier programming of more complex tasks.

The wiring diagram for this setup is presented in **Supplementary Figure 1**. Output from the REM indicating rotation and direction was sent to 1) a data collection PC via USB, 2) a neural data acquisition system via TTL pulses, and 3) to the ECM. The ECM performed multiple tasks such as calculating pull speed and distance and delivering reward contingent on distance pulled and choices made by the animal. Software for the REM and ECM is provided at https://github.com/CowenLab/String_Pulling_System.

### *String pulling in unilateral 6-OHDA-lesioned parkinsonian rats:*

To further test the translatability of the string-pulling task and the novel automated apparatus in a Parkinson’s disease (PD) model, we first established the use with PD animals. We verified the usefulness of the string-pulling task in this PD model, and the Results from manual blinded investigator scoring are shown in **Supplementary Figure 2** with semi-quantitative western verification of the lesion provided in **Supplementary Figure 3.** For the investigator scoring experiment animals were habituated to the experimenter, reward (cheerios), and apparatus (26 cm x 26 cm with a transparent window) for 3 days. Day one of training consisted of two trials with 10 strings of varying lengths (half rewarded). In day two, a single 100 cm string was baited and hung in front of the apparatus. The trial ended when the rat retrieved the reward or 5 minutes elapsed. Once a rat successfully retrieved the cheerio in 8 consecutive trials, the string length was increased to 150 cm. Training was complete once a rat retrieved the reward in 8 trials with the 150 cm string. Pre-LX testing consisted of the same protocol as the last day of training. Following Pre-LX testing, the rats (n = 3/group) received either a unilateral 6-OHDA lesion (>90%) or sham-surgery in the medial forebrain bundle as previously established [1]. The videos were scored by blinded experimenters and the data analyzed with GraphPad Prism 8.0 software (GraphPad Software, La Jolla, CA).

To illustrate the potential of the novel automated system we then conducted an experiment using the automated string-pulling apparatus in a single animal (**Supplementary Figure 4**).

### *Tyrosine hydroxylase western blot analysis:*

Confirmation of the 6-OHDA lesion in PD rats shown in Supplementary Figure 2 was confirmed using western blot analysis as published in Bartlett et al. [36] 8 μg of striatal protein from each hemisphere was loaded for gel electrophoresis. Gels were then transferred and incubated with primary antibodies: Tyrosine Hydroxylase (TH) (1:2000; AB152, Millipore, Billerica, MA), Beta-Actin (1:10,000; A1978, Sigma-Aldrich, St. Louis, MO), followed by secondary antibodies: IRDye® 800 CW Donkey anti-Rabbit (1:20,000; LI-COR Biosciences, Lincoln, NE), IRDye® 680 CW Donkey anti-Mouse (1:20,000; LI-COR

Biosciences). Blots were scanned on an Odyssey CLx imager and analyzed with Empiria Studio Software (LI-COR Biosciences).

### *Near-Infrared Tyrosine Hydroxylase Immunohistochemistry and Densitometry analysis:*

Confirmation of the 6-OHDA lesion in the single 6-OHDA treated rat shown in Supplementary Figure 3 was confirmed using near-infrared imaging. Three coronal sections containing the striatum were selected at AP: -0.4, -0.2, and +0.2, relative to bregma, according to the atlas of Paxinos and Watson. [45] Sections were incubated in a blocking buffer containing Odyssey Intercept® (TBS) Blocking buffer (LI-COR Biosciences) and 10% normal donkey serum (Jackson ImmunoResearch Laboratories Inc., West Grove, PA) at room temperature for one hour. Sections were then incubated in blocking buffer, with the addition of 0.1% Tween-20, for TH (1:2000; Millipore) overnight at 4 °C. Tissue was then washed 3 x 5 min in TBS + 0.1% Tween-20 (TBST) and incubated in blocking buffer and 0.1% Tween-20 including the IRDye® 800 CW Donkey anti-Rabbit (1:200; LI-COR Biosciences) secondary antibody for one hour at room temperature. Sections were washed again 3 x 5 minutes in TBST, mounted on subbed slides and imaged on an Odyssey CLx imager (800 nm channel, 42 μm resolution; LI-COR Biosciences). TH-immunoreactivity was determined as previously published in Yue et al. [46] using Image Studio Lite (LI-COR Biosciences) to quantify signal intensity.

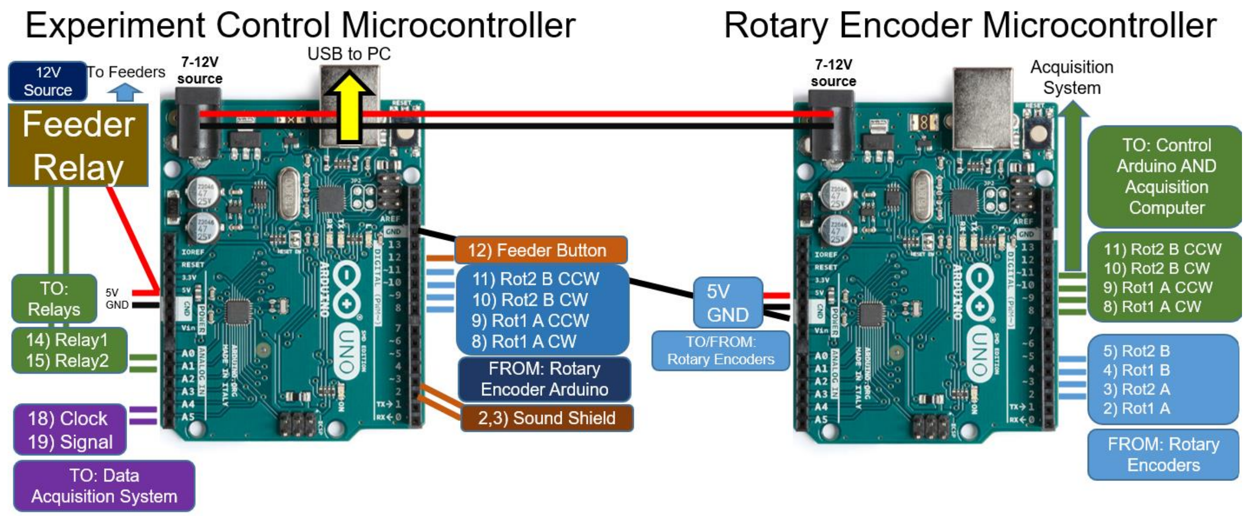


**Supplementary Figure 1** Pinout and wiring diagram of the dual-microcontroller setup. This setup reduces the computational demands and simplifies programming of the experiment control microcontroller by separating processing of rotary encoder measurement from experiment control.

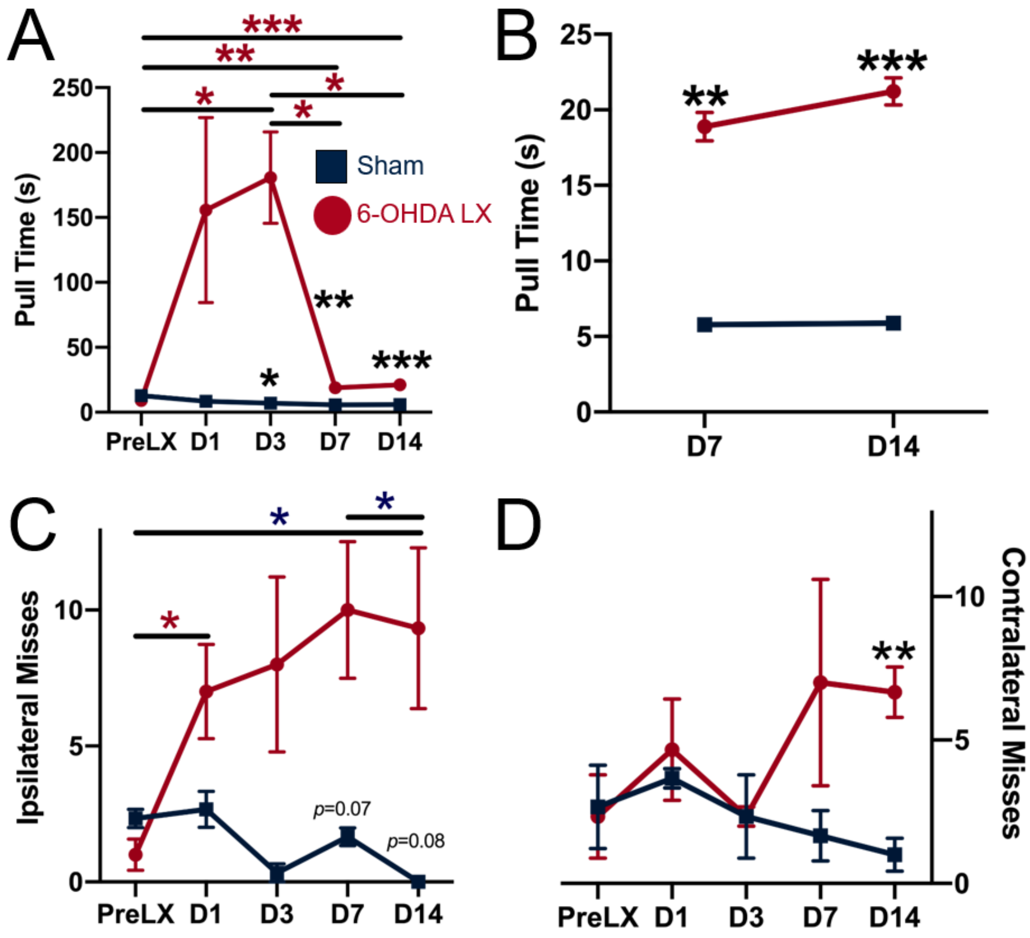


**Supplementary Figure 2:** A 6-OHDA lesion reduces the latency to complete the string-pulling task. A) the mean latency ±SEM to complete the task for each testing session. PreLx: pre 6-OHDA lesion session. D1-D14 are days post-lesion. N = 3/group, *p<0.05, ** p<0.01, *** p<0.001, RM One-way ANOVA, Fisher's LSD post-hoc test (red asterisks). B) Zoomed in image of the D7 and D14 days in (A) to emphasize the effect on pull behavior once the lesion is complete. C) Mean number of ipsilateral misses ±SEM in each testing session. There is a significant decrease in ipsilateral misses in sham rats by Day 14, as compared to pre-lesion. On Day 7 and 14, 6-OHDA-lesioned rats show a non-significant increase (p = 0.07 and p = 0.08 vs. sham-lesion) in ipsilateral misses compared to sham rats. D) 6-OHDA-lesioned rats demonstrate a significant increase in contralateral misses by Day 14. n = 3/group, *p < 0.05, **p < 0.01, RM One-way ANOVA, Fisher's LSD post-hoc tests (red and blue asterisks); Two-way ANOVA, Fisher's LSD post-hoc tests (black asterisks).

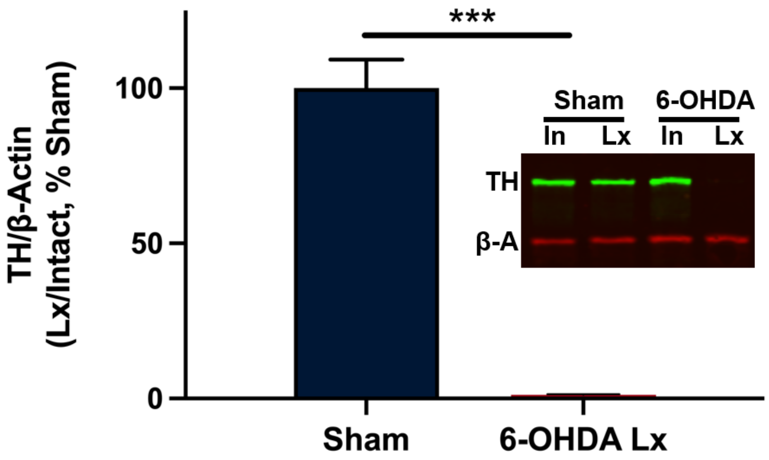


**Supplementary Figure 3.** Semi-quantitative western verification of unilateral 6-OHDA lesion from the rats (n = 3/group) depicted in Supplementary Figure 2. The graph shows the quantification of tyrosine hydroxylase (TH) with sham *vs*. 6-OHDA treated rats plotted as % loss (mean ± SEM). Two-tailed *t*-test, ***$p < 0.001$. Inset shows example TH (green) and beta-actin (red) in the intact and lesioned side of sham and 6-OHDA treated rats.

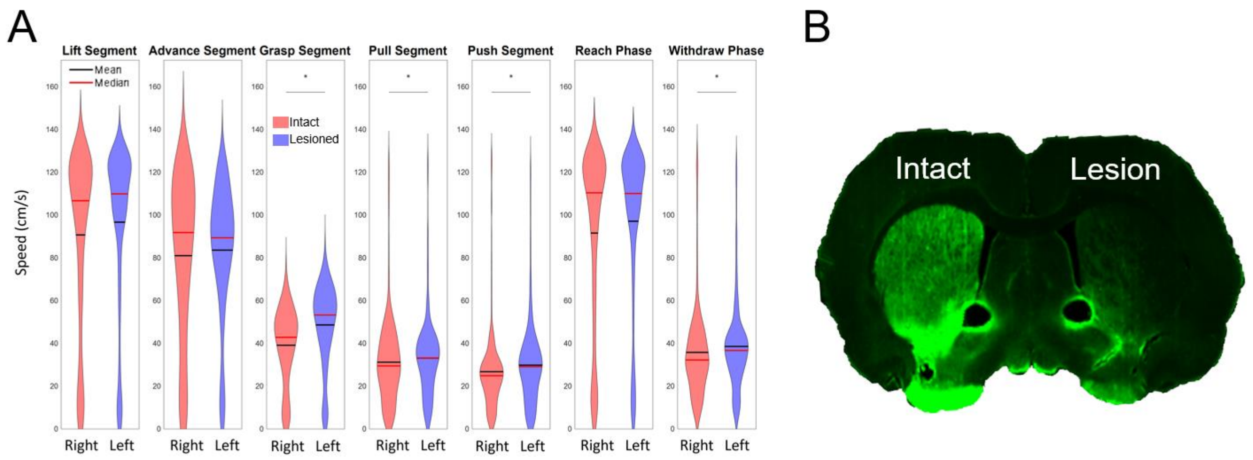


**Supplementary Figure 4:** A) Speed (cm/sec) of the left and right paw recorded from a 6-OHDA hemi-lesioned PD model animal (from 1 rat) with a right hemisphere lesion. Data shown for each phase of the pulling behavior as well as the general response to the Reach and Withdrawal phases. Lines and asterisks indicate statistically significant differences ($p < 0.05$ with Holm adjustment for multiple comparisons, Wilcoxson test) in the distributions of peak speeds of the left and right paws. We predicted that the speed of the affected limb (left paw) would be lower than the right paw. Instead, peak speed during the grasp, pull, and push segments and withdrawal phase were faster for the left relative to the right paw. These data illustrate the application of the system for assessing behavior in animal models of disease, but given the sample size, these data are only illustrative. B) Verification of unilateral 6-OHDA lesion from the rat (n = 1) depicted in A. Image shows TH-immunoreactivity (green) in the intact and lesioned striatum of the rat unilaterally injected with 6-OHDA.

| Table 1 - Parts List | | | |
|---|---|---|---|
| Component | Manufacturer | Part No. | Website |
| Electronic Components | | | |
| Mako U-130b Camera* (option1) | Allied vision | Mako U-130b | https://cdn.alliedvision.com/fileadmin/pdf/en/Mako_U-130_DataSheet_en.pdf |
| Alvium 1800 U-040 (option 2) | Allied vision | Alvium 1800 U-040 | https://www.alliedvision.com/en/products/alvium-configurator/alvium-1800-u/040-1/#_configurator |
| LED Panel | Viltrox | L116T | https://www.amazon.com/VILTROX-L116T-3300K-5600K-Temperature-Brightness/dp/B07D8TTFSR/ref=sr_1_3?dchild=1&keywords=viltrox+led&qid=1620409459&sr=8-3 |
| Ring Light, 10" | UBeeSize | UBeeSize Ring Light | https://www.amazon.com/10-Extendable-UBeesize-Ringlight-Compatible/dp/B07QFV72LK/ref=sr_1_3?dchild=1&keywords=ubeesize+ring+light&qid=1620409432&sr=8-3 |
| Arduino Uno | Arduino | Arduino Uno Rev3 | https://store-usa.arduino.cc/collections/boards/products/arduino-uno-rev3 |
| Arduino Mega | Arduino | Mega 2560 Rev3 | https://store.arduino.cc/usa/mega-2560-r3 |
| Rotary Encoder | BQLZR | BQLZR 600P/R | https://www.amazon.com/gp/product/B00UTIFCVA/ref=ppx_yo_dt_b_search_asin_title?ie=UTF8&psc=1 |
| Digital Distance Sensor 5cm | Pololu | 4050 | https://www.pololu.com/product/4050 |
| Project Box 12.2 x 11.2 x 4.5" | Zulkit | NA | https://www.amazon.com/gp/product/B08NGGMKC3/ref=ppx_yo_dt_b_search_asin_image?ie=UTF8&th=1 |
| Solenoid Valve 1/4" | STC Valve | 2P025-1/4 | https://www.stcvalve.com/Solenoid-Valve-Specifications-2P025-Series.htm |
| Panel Mount Aviation Connectors | Hilitchi | 8541770567 | https://www.amazon.com/gp/product/B07F5B5LLX/ref=ppx_yo_dt_b_search_asin_title?ie=UTF8&th=1 |
| Physical Apparatus | | | |
| T-slot nuts | Sutemribor | STBR-T-luomu-160P-kit | https://www.amazon.com/gp/product/B07FPLZXTF/ref=ppx_yo_dt_b_search_asin_image?ie=UTF8&psc=1 |
| Cotton Twine #16 [CHECK] | Ace Hardware | C8016B0008AC | https://www.acehardware.com/departments/hardware/chain-and-rope/ropes/7373707 |
| 3D printer wheel with bearings | SeekLiny | NA | https://www.amazon.com/dp/B09SSWZG82?ref=ppx_yo2ov_dt_b_product_details&th=1 |
| Extruded Aluminum (20/20) | Zyltech | EXT-2020-REG-1000-10X | https://www.amazon.com/gp/product/B07F6R2Z99/ref=ppx_yo_dt_b_search_asin_title?ie=UTF8&psc=1 |
| 90° Corner Bracket | LANIAKEA | NA | https://www.amazon.com/gp/product/B08NXGV1YH/ref=ppx_yo_dt_b_search_asin_title?ie=UTF8&psc=1 |
| Neural Data Acquisition Systems | | | |
| Intan USB Interface Board (option 1)* | Intan Technologies | | https://intantech.com/RHD_USB_interface_board.html |
| RHD Recording Controller (option 2) | Intan Technologies | | https://intantech.com/recording_controller.html |
| Neuropixels recording system | Imec | | https://www.neuropixels.org/ |
| * Model Discontinued. The Alvium 1800 is the updated model. | | | |

**Supplementary Table 1:** Parts list with hyperlinks
